# Genetic mapping with Allele Dosage Information in Tetraploid *Urochloa decumbens* (Stapf) R.D. Webster Reveals Insights into Spittlebug (*Notozulia entreriana* Berg) Resistance

**DOI:** 10.1101/360594

**Authors:** Rebecca Caroline Ulbricht Ferreira, Letícia Aparecida de Castro Lara, Lucimara Chiari, Sanzio Carvalho Lima Barrios, Cacilda Borges do Valle, José Raul Valério, Fabrícia Zimermann Vilela Torres, Antonio Augusto Franco Garcia, Anete Pereira de Souza

## Abstract

*Urochloa decumbens* (Stapf) R.D. Webster is one of the most important African forage grasses in Brazilian beef production. Currently available genetic-genomic resources for this species are restricted mainly due to polyploidy and apomixis. Therefore, crucial genomic-molecular studies such as the construction of genetic maps and the mapping of quantitative trait loci (QTLs) are very challenging and consequently affect the advancement of molecular breeding. The objectives of this work were to (i) construct an integrated *U. decumbens* genetic map for a full-sibling progeny using GBS-based markers with allele dosage information, (ii) detect QTLs for spittlebug (*Notozulia entreriana*) resistance, and (iii) seek putative candidate genes involved in resistance/defense against pathogens. We used the *Setaria viridis* genome as reference to align GBS reads and selected 4,240 high-quality SNPs markers with allele dosage information. Of these markers, 1,000 were distributed throughout nine homologous groups with a cumulative map length of 1,335.09 cM and an average marker density of 1.33 cM. We detected QTLs for resistance to spittlebug, an important pasture insect pest, that explained between 4.66% and 6.24% of the phenotypic variation. These QTLs are in regions containing putative candidate genes related to resistance/defense against pathogens. Because this is the first genetic map with SNP autotetraploid dosage data and QTL detection in *U. decumbens*, it will be useful for future evolutionary studies, genome assembly, and other QTL analyses in *Urochloa* spp. Moreover, the results might facilitate the isolation of spittlebug-related candidate genes and help clarify the mechanism of spittlebug resistance. These approaches will improve selection efficiency and accuracy in *U. decumbens* molecular breeding and shorten the breeding cycle.

## 1 Introduction

Brazilian cultivated pastures are the basis for the production of beef cattle, and they cover extensive areas, most of which are populated by grasses of the genus *Urochloa* (syn. *Brachiaria*) (Jank et al., 2011; Associação Brasileira das Indústrias Exportadoras de Carne (ABIEC), 2016). This genus belongs to the Poaceae family and comprises species of plants distributed in tropical and subtropical regions, mainly in the African continent (Renvoize et al., 1996; Valle et al., 2009).

One of the most widely cultivated species of genus is *Urochloa decumbens* (Stapf) R.D. Webster, also known as signalgrass. This forage grass has relevant agronomic attributes, including exceptional adaptation to poor and acidic soils that are typical of the tropics, leading to good animal performance (Valle et al., 2010). However, the species is susceptible to several types of spittlebug, including *Notozulia entreriana* Berg (Hemiptera: Cercopidae), which is the most damaging pest for tropical pastures (Gusmão et al., 2016).

Spittlebugs reduce biomass production, decrease palatability, and reduce the carrying capacity of pastures (Valério and Nakano, 1988). Therefore, breeding programs aim to develop new cultivars with reasonable or high resistance to spittlebugs to reduce forage loss. To achieve this advance, it is essential to understand the genetic mechanisms involved in the spittlebug resistance response in forage grass and to identify valuable markers for selection in breeding schemes. Currently, spittlebug resistance in *Urochloa* grasses is thought to be under relatively simple genetic control and is highly heritable (Miles et al., 1995). Therefore, despite being a trait with little genetic information, spittlebug resistance can probably be easily manipulated in a plant breeding program.

Genetic breeding of *U. decumbens* is recent and has proven challenging because this grass is predominantly tetraploid (2n = 4x = 36) and apomictic (Naumova et al., 1999; Valle et al., 2008). In this scenario, the number of currently available molecular genetic resources is limited. Previous molecular studies have increased knowledge of *U. decumbens* genetics, including the development of sets of microsatellite markers (Ferreira et al., 2016; Souza et al., 2018) and the first transcriptome (Salgado et al., 2017) of the species as well as linkage maps from the interspecific progeny of *U. decumbens* × *U. ruziziensis* (Worthington et al., 2016). Other studies with molecular markers have analyzed the genetic relationships of this species to other species in the *Urochloa* genus (Almeida et al., 2011; Pessoa-Filho et al., 2017; Triviño et al., 2017).

Until recently, it was not possible to perform intraspecific crossings due to the ploidy barriers between apomictic (polyploid) and sexual (diploid) accessions, but this restriction changed when diploid accessions were artificially tetraploidized using colchicine (Simioni and Valle, 2009). This advance allowed the generation of a base population and the exploration of a genetic variability previously conserved by apomixis, favoring new molecular studies, including building specific genetic maps to increase the genetic knowledge of the species.

Genetic maps are essential tools for genetics and genomics research, and they are becoming increasingly common in polyploids species, mainly due to the development of next-generation-sequencing (NGS) technologies and advances in genetic and statistical methods (Garcia et al., 2013). In this study, we utilized the genotyping-by-sequencing (GBS) technique, which enables the discovery of thousands of SNPs at low cost (Elshire et al., 2011; Poland et al., 2012). In addition, robust software has been developed for polyploid organisms, including programs that can estimate the allele dosage of SNPs and provide more genetic information to generate linkage maps with high marker densities (Serang et al., 2012; Garcia et al., 2013; Mollinari and Serang, 2015; Bourke et al., 2016, 2018).

The mapping process is more complex in polyploid species, mainly because of the larger number of possible genotypes (Ripol et al., 1999; Vigna et al., 2016). Most polyploid genetic maps have been limited to the use of single-dose markers (SDMs). In this case, either each marker is considered as a single-allele copy from only one of the parents of the cross, with a 1:1 segregation ratio, or the SDMs in both parents segregate in a 3:1 ratio (Wu et al., 1992). This method has been developed and successfully adopted for various forage grass species, such as *Panicum maximum* (Ebina et al., 2005), *Paspalum notatum* (Stein et al., 2007), *Panicum virgatum* (Okada et al., 2010) and *Urochloa humidicola* (Vigna et al., 2016). However, the use of only SDMs represents a loss of genetic information, because in autotetraploid species, for example, five possible dosage classes exist: nulliplex (aaaa), simplex (Aaaa), duplex (AAaa), triplex (AAAa), and quadruplex (AAAA). Thus, many other marker segregations can be considered. Knowledge of the dosage of an SNP is essential for genetic studies in polyploids species and can significantly increase the information imparted by each locus (Voorrips et al., 2011; Garcia et al., 2013; Bourke et al., 2016, 2018).

High-resolution genetic linkage mapping for polyploid species can identify beneficial trait loci and allow genomics-based breeding programs (Shirasawa et al., 2017). Molecular markers have been widely used to locate quantitative trait loci (QTL) associated with quantitative resistance to insects in many crop plants; however, genomic loci related to resistance to spittlebugs have not been determined in any plant species. Particularly in *U. decumbens*, the identification of loci involved in this trait is a promising tool for characterizing the genetic architecture and can assist in the design of strategies to be introduced into breeding programs to increase the efficiency of the selection processes and accelerate the release of new cultivars (Valle et al., 2009).

This study reports the development and application of GBS for mapping studies in an intraspecific progeny of *U. decumbens*. To the best of our knowledge, there have been no reports of QTL mapping in signalgrass. Our goals were to (i) build a GBS-based integrated genetic map using autotetraploid allele dosage information in a bi-parental progeny, (ii) identify QTLs related to spittlebug resistance on the integrated genetic map, and (iii) search for putative candidate genes that may be involved in spittlebug resistance.

## 2 Materials and Methods

### 2.1 Plant Material and DNA Extraction

At the Brazilian Agricultural Research Corporation (Embrapa) Beef Cattle (EBC), Campo Grande/MS, an intraspecific cross was made between *U. decumbens* D24/27 (sexual accession tetraploidized by colchicine) and *U. decumbens* cv. Basilisk (tetraploid, apomictic cultivar used as the pollen donor); therefore, both parents were tetraploid. The full-sib progeny of 239 F_1_ individuals was analyzed, from which 217 hybrids were identified using SSRs markers obtained from Ferreira et al. (2016).

Leaf samples from each hybrid and both parents were collected, and genomic DNA was extracted using the DNeasy 96 Plant Kit (Qiagen GMbH, Germany). DNA concentrations were determined using a Qubit fluorometer (Invitrogen, Carlsbad, CA).

### 2.2 GBS Library Construction and Sequencing

The GBS library was prepared at the University of Campinas following the protocol described by Elshire et al. (2011). Samples from both parents of the progeny were replicated five times for sequencing. Each individual within a library was part of a 96-plex reaction. Each DNA sample (300 ng in a volume of 10 μL) was digested with the restriction enzyme NsiI (NEB) to reduce genomic complexity and then ligated to a unique barcoded adapter plus a common adapter. The 96-plex libraries were checked for quality using an Agilent DNA 1000 Kit on an Agilent 2100 Bioanalyzer (Agilent Technologies, Santa Clara, CA, United States). Libraries sequencing was performed as 150-bp single-end reads on the Illumina NextSeq 500 platform. The quality of the resulting sequence data was then evaluated with the NGS QC Toolkit v2.3.3 (Patel and Jain, 2012).

### 2.3 SNP Calling

SNP discovery and genotype calling were performed using Tassel-GBS pipeline (Glaubitz et al., 2014), which was modified to obtain the original count of the number of reads for each SNP allele (Pereira et al., submitted). Because this pipeline requires a reference genome, and because the *U. decumbens* genome has not yet been sequenced, we aligned the GBS tags using the *Setaria viridis* genome (v 1.0; ~394.9 Mb arranged in 9 chromosomes and 724 scaffolds; diploid forage) as an alternative pseudo-genome. This genome is available from the Phytozome website (http://www.phytozome.net/) (Goodstein et al., 2012), and it sequence data were produced by the US Department of Energy Joint Genome Institute. The Bowtie2 algorithm version 2.1 (Langmead and Salzberg, 2012) was used to align tags against the pseudo-reference with -D and -R parameters defined as 20 and 4, respectively, and with the very-sensitive-local argument.

### 2.4 Allele Dosage Estimation and Data Filtering

SuperMASSA software (Serang et al., 2012; Pereira et al., submitted) was used to estimate the allele dosage (number of copies of each allele) of each individual. The minimum overall depth considered was 25 reads, and the model used was the F_1_ Population Model. Markers were fitted and filtered to ploidy 4.

Additionally, monomorphic SNPs were removed, and markers with a maximum of 25% missing data were selected manually using R software.

### 2.5 Linkage Map

An integrated linkage map was built using the TetraploidSNPMap (TPM, BioSS) software (Hackett et al., 2017), following the methodology described in Hackett et al. (2013, 2014). Homologous groups were constructed separately using the SNPs identified on each relative *S. viridis* chromosome. Markers with significance of the χ^2^ goodness-of-fit statistic less than 0.001 for simplex and 0.01 for higher segregations were considered distorted and were removed. A total of 2,725 distorted SNPs were excluded from the cluster analyses. The remaining 1,515 non-distorted SNPs were clustered into 9 homologous groups using a Chi-square test for independent segregation (Hackett et al., 2013).

Considering the ordination of markers, a 2-point analysis was used to calculate the recombination frequency and LOD score for each pair of SNPs in each possible phase by an expectation-maximization (EM) algorithm to maximize the likelihood. Duplicate and near-duplicate SNPs were removed in this step. The recombination frequencies were converted to map distances using Haldane’s mapping function, and a multi-dimensional scaling analysis (MDS) was then performed to calculate the best order for the SNPs in the homology group. After this step, we excluded some SNPs that showed low LOD scores or that were distant from the rest of the group (Hackett et al., 2017).

Some phases of the ordered SNPs were inferred by the automated phase analysis in TetraploidSNPMap, and others were completed manually based on the most likely phase for previous informative pairs prior to carrying out QTL analysis.

The genetic map was drawn using MapChart 2.32 (Voorrips, 2002), in which SNPs configurations were identified with different colors.

### 2.6 Phenotypic Evaluation

The resistance of *U. decumbens* to spittlebugs (*N. entreriana*) was evaluated in greenhouse experiments at Embrapa Beef Cattle, Brazil, according to the methodology described by Lapointe et al. (1992) and Valério et al. (1997). From 2011 to 2013, a total of 12 experiments were conducted; each experiment had a randomized complete block design (RCBD) with different numbers of blocks and treatments. Common treatments were repeated in all experiments and were used as checks in the statistical analysis. A total of 349 individuals were evaluated for spittlebug resistance, of which 157 were hybrids of the mapping population.

During 2011 and 2012, five RCBD experiments were conducted, with ten blocks and 114 individuals evaluated. The best hybrids were selected with respect to spittlebug resistance and included in the experiments conducted in 2012 and 2013, when seven RCBD experiments were conducted, with four blocks and 259 individuals. The common checks for all experiments were *U. decumbens* cv. Basilisk, *U. brizantha* cv. Marandu, *Urochloa* spp. cv. BRS Ipyporã, and *Urochloa* spp. MulatoII. The variable evaluated was the percentage of nymphal survival (number of nymphs that reached the adult stage, as a percentage). The following statistical linear mixed model was used for the analysis:

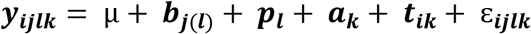

where *y*_*ijlk*_ is the phenotype of the *i*-th individual, at *j*-th block, *l*-th experiment, and *k*-th year; μ is the overall mean; *b*_*j*(*l*)_ is the random effect of the *j*-th block within the *l*-th experiment 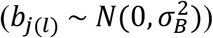; *p*_*l*_ is the random effect of the *l*-th experiment 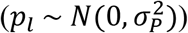; *a*_*k*_ is the fixed effect of the *k*-th year; *t*_*ik*_ is the random effect of the *i*-th individual in the *k*-th year (*t*_*ik*_ ~ *N*(0, **G**)), in which ***G*** is the variance-covariance (VCOV) matrix for genetic effects; and ɛ_*ijlk*_ is the residual error 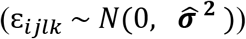. The ***G*** matrix is indexed by two factors (genotype and year) written as the Kronecker product of matrices, ***G*** = ***l***_***n***_ ⊗ ***G***_***a***_, in which ***I***_***n***_ is an identity matrix relative to genotype effects and ***G***_***a***_ is the VCOV matrix relative to year effects. The ***G***_*a*_ matrix was evaluated considering six different VCOV structures: independent (ID), diagonal (DIAG), compound symmetry (CS), compound symmetry heterogeneous (CS-Het), first-order factor analytic (FA1), and unstructured (US). The selection model was performed considering the Akaike Information Criteria (AIC) (Akaike, 1974) and Schwarz Information Criteria (SIC) (Schwarz, 1978).

Phenotypic analysis was performed using the packages ASReml-R v3 (Butler et al., 2009; Gilmour et al., 2009) and ASRemlPlus (Brien, 2016). Heritability between genotype means was estimated using the index as presented by Cullis et al. (2006) and Piepho and Möhring (2007), where:

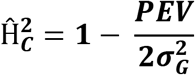
in which *PEV* is the prediction error variance, and 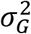 is the genetic variance.

### 2.7 QTL Mapping

QTL mapping for spittlebug resistance was performed by applying an Interval Mapping (IM) model based on autotetraploid allele dosage information with TetraploidSNPMap software (Hackett et al., 2017). This analysis was performed separately for each homology group using the SNP data for each homology group, the integrated genetic map with phase information and the phenotypic trait data.

The interval mapping was fit to a model of additive effects using a weighted regression approach. A significant QTL was declared if the LOD score was above the 95% threshold obtained from a permutation test with 1,000 permutations. From a significant QTL, simple models such as a simplex, duplex or double-simplex QTL models were then tested using the SIC (Schwarz, 1978) to determine if the estimated genotype corresponded to the most likely QTL location. The SIC is calculated in TetraploidSNPMap as follows:

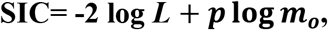

where *L* is the likelihood for the simple model, *p* is the number of parameters in the simple model, and *m*_o_ is the number of observations (the 36 genotype means). Models with the lowest value for the SIC were considered the best models (Hackett et al., 2014).

If two closely linked significant QTLs were identified on the same chromosome, TetraploidSNPMap could not estimate the individual QTL LOD and parameter values, yielding data only pertaining to the largest effect.

### 2.8 Search for Candidate Genes in Detected QTL Regions

We investigated the functional annotation of the candidate genes located close to the QTL regions associated with spittlebug resistance. Using the chromosomal locations of the markers adjacent to QTLs in the *S. viridis* genome as a reference, we aligned the sequences found in these regions (100 kb approximately) with a BLASTX search (e-value cutoff of 1e-0.5) against plants databases using the JBrowse tool (http://www.phytozome.org/). Finally, from this alignment, we selected only genes reported in the literature as associated with resistance/defense against pathogens that were present in the transcriptome of *U. decumbens* (Salgado et al., 2017), our species of study.

## 3 Results

### 3.1 SNP Calling

The GBS library generated a total of 1,183,089,925 raw reads (94% Q20 bases), with an average of 4,151,193 reads per individual. Using the Tassel-GBS pipeline modified to polyploid, alignment with the *S. viridis* genome enabled the identification of 58,370 SNPs markers. The output of the Tassel-GBS pipeline was used as the input for the SuperMASSA software, and 9,345 SNPs markers with allele dosage were selected with a minimum overall depth of 25 reads. Afterward, markers with up to 25% missing data and monomorphic markers were removed, resulting in 4,240 SNPs markers that were used in the subsequent steps.

Using three different SNPs markers, Figure 1 shows how the SuperMASSA software uses the ratio of allele count to classify individuals according to their genotypes using a probabilistic graphical model (Serang et al., 2012). The markers were named with the allele dosage segregation, the homology group number, and the chromosomal position of the SNP in *S. viridis* genome.

**Figure 1.**
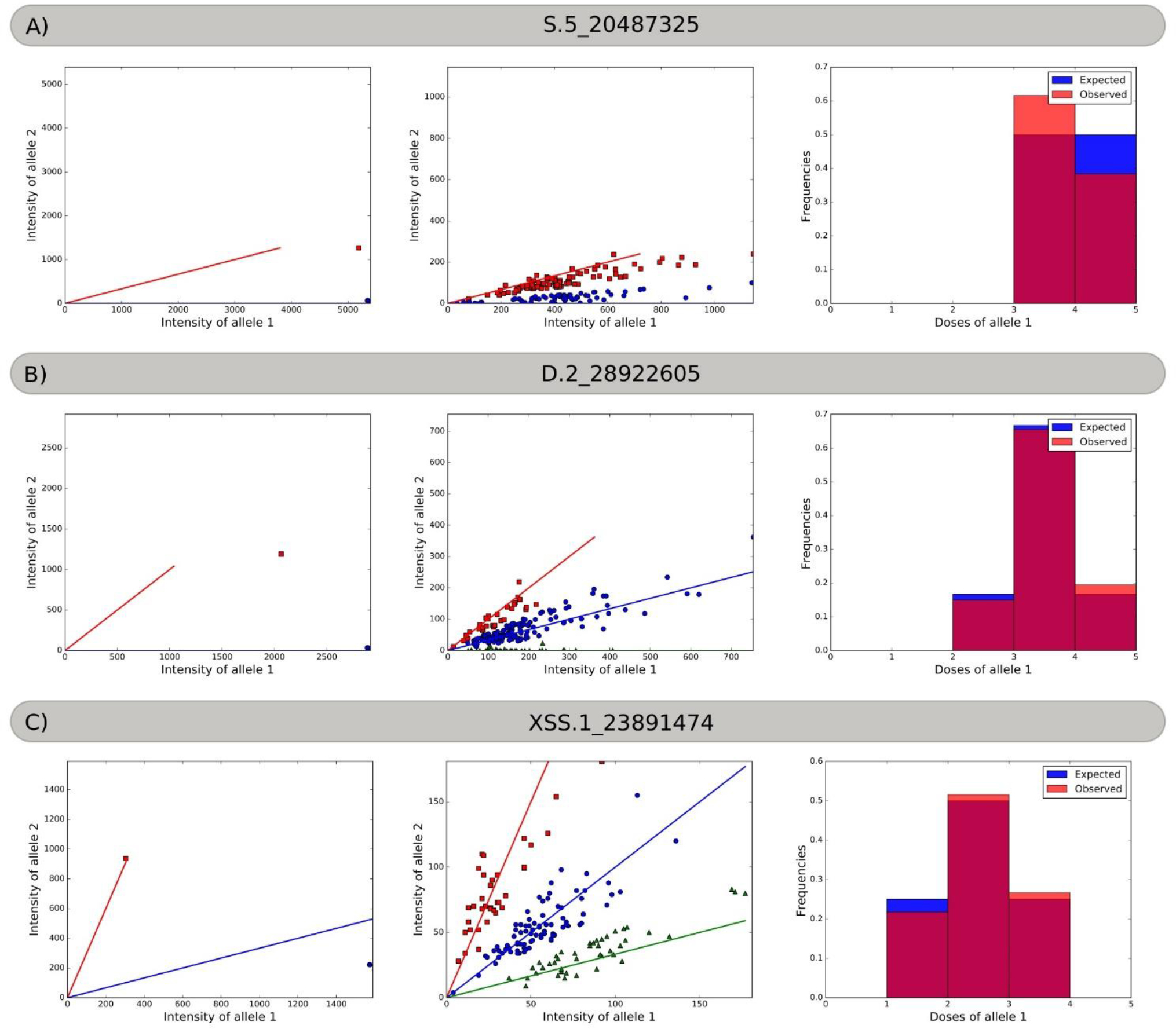
Allele dosage in the parents and progeny, and the frequency histograms. A) Marker S.5_20487325. Red squares represent the Aaaa parent and offspring, and blue circles represent the aaaa parent and offspring. B) Marker D.2_28922605. Red squares represent the AAaa parent and offspring, blue circles represent the aaaa parent and the Aaaa offspring, and green triangles represent the aaaa offspring. C) Marker XSS.1_23891474. Red squares represent AAAa parent and offspring, blue circles represent the Aaaa parent and the AAaa offspring, and green triangles represent the Aaaa offspring.

**Figure 2.**
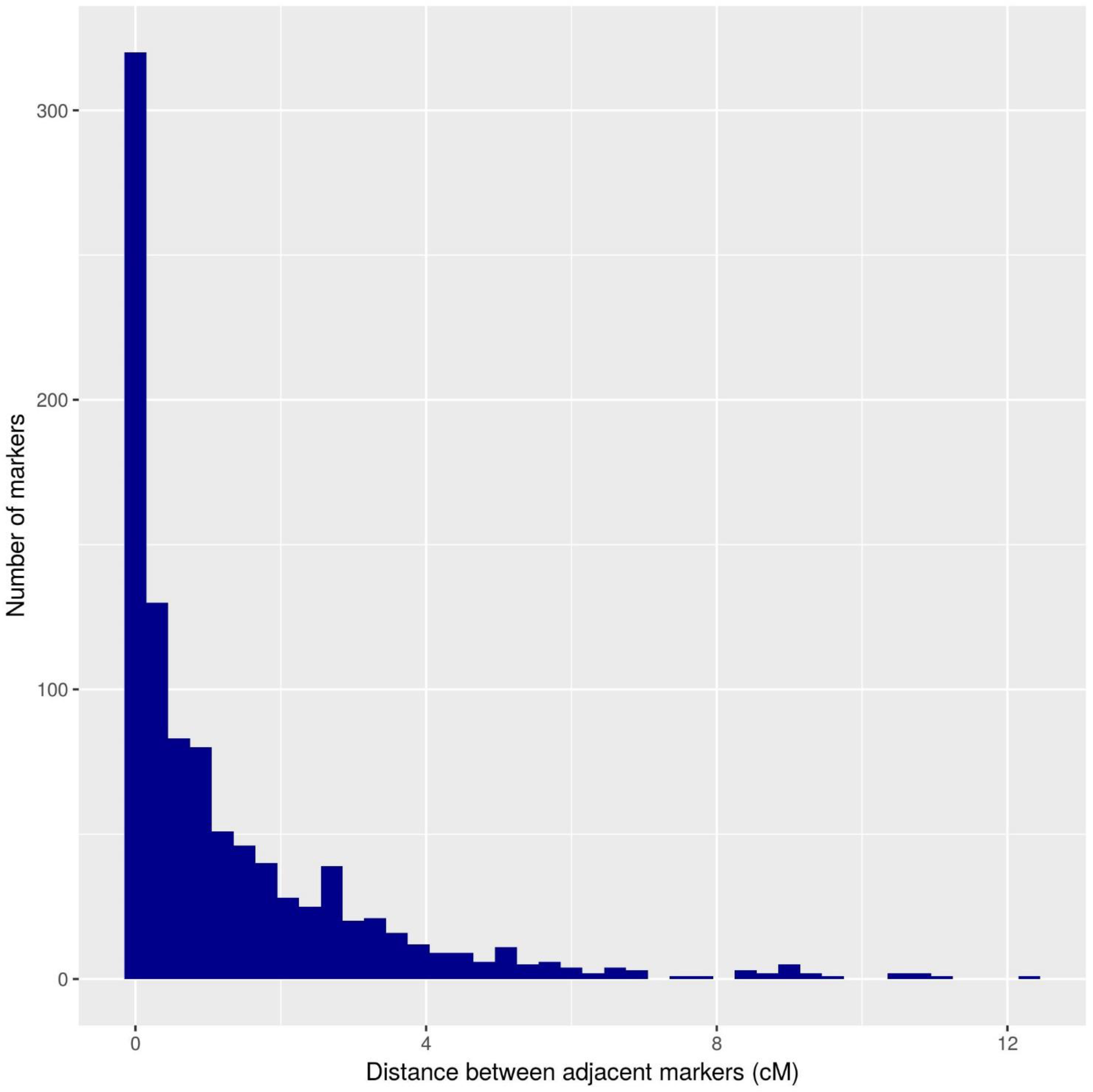
Histogram of the distance between adjacent markers on the *U. decumbens* genetic map.

GBS sequence data has been submitted to the NCBI Sequence Read Archive (SRA) under accession number SRP092493.

### 3.2 Genetic Map

An integrated genetic map was built using 217 full sibs obtained from a cross between *U. decumbens* cv. Basilisk (apomictic cultivar) and *U. decumbens* D24/27 (sexual accession). Of the 4,240 markers used for linkage analysis, 1,515 showed significance of the χ^2^ goodness-of-fit statistic, and 1,000 were placed in the linkage map (Table 1).

**Table 1.** The number of GBS-based markers after significance testing, the number of mapped markers within each homology group (HG), and the length and marker density of each HG (cM) in the genetic map.

| HG | No. GBS-based markers* | No. mapped markers | Length of HG (cM) | Marker density (cM) |
| --- | --- | --- | --- | --- |
| 1 | 132 | 100 | 136.74 | 1.37 |
| 2 | 181 | 121 | 164.95 | 1.36 |
| 3 | 219 | 138 | 151.56 | 1.10 |
| 4 | 169 | 101 | 139.70 | 1.38 |
| 5 | 183 | 123 | 129.70 | 1.05 |
| 6 | 100 | 63 | 161.72 | 2.57 |
| 7 | 154 | 99 | 140.08 | 1.41 |
| 8 | 82 | 68 | 144.07 | 2.12 |
| 9 | 295 | 187 | 166.57 | 0.89 |
| Total | 1,515 | 1,000 | 1,335.09 | 1.33 |

\*Markers were selected if the $\chi^2$ goodness-of-fit statistic was higher than 0.001 for simplex and 0.01 for higher segregations.

The markers were distributed throughout nine homology groups (HGs), with a cumulative map length of 1,335.09 cM and an average marker density of 1.33 cM. The length of each group ranged from 129.70 cM (HG7) to 166.57 cM (HG9), with an average of 148 cM (Table 1). Homology group 9 showed the highest average density and the highest number of mapped markers (Table 1; Figure 3).

**Figure 3.**
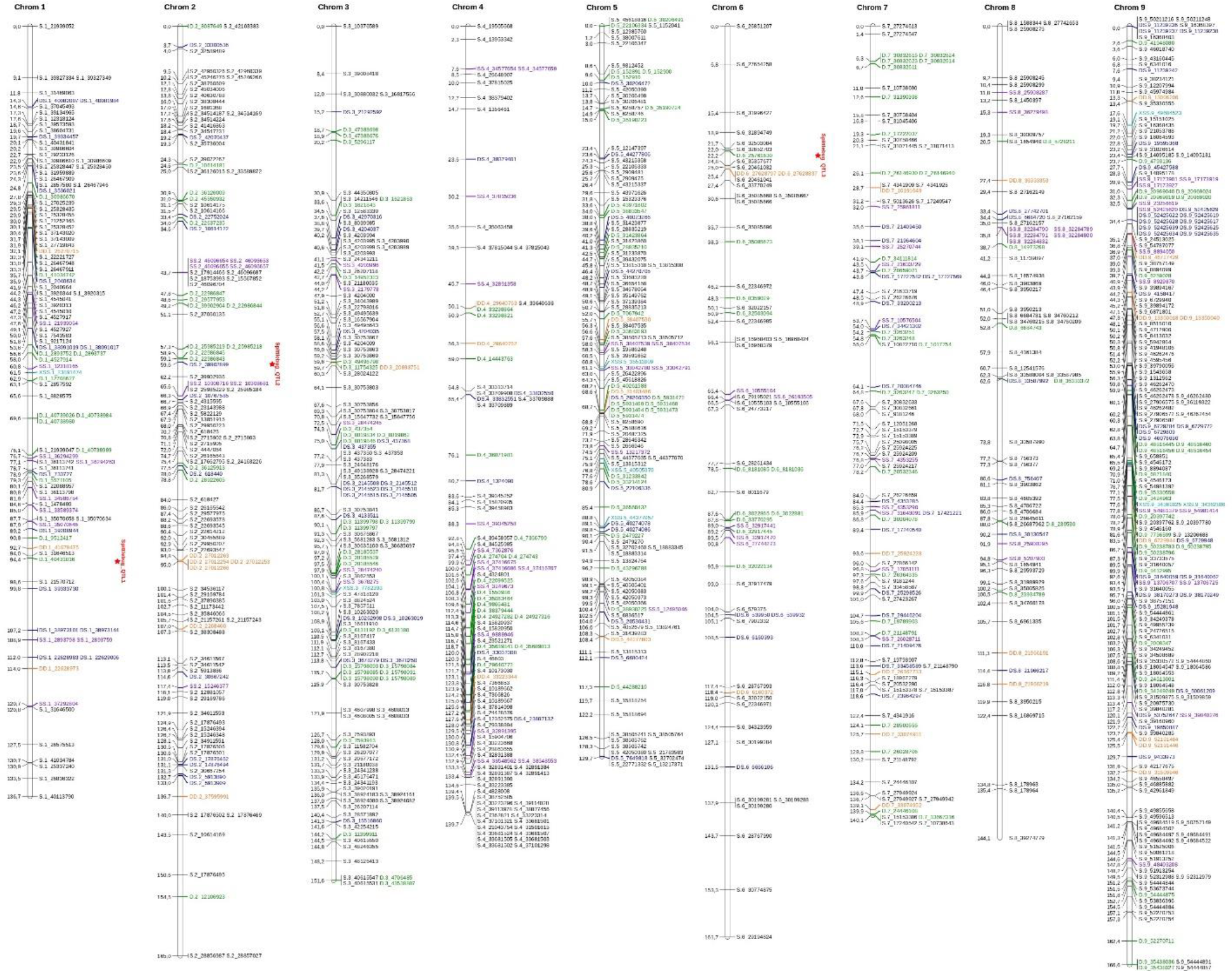
Linkage map for *U. decumbens*: homologous groups from 1 to 9. The genotype configuration of each marker is indicated by the marker name prefix and color (Table 2). The QTLs are identified in HG1, HG2 and HG6.

The distances between adjacent markers on the genetic map were plotted as a histogram, which revealed that the majority of markers were within 1 cM of each other (Figure 2). The two largest distances between adjacent markers were located in HG8 and had lengths of 11.2 cM and 12.4 cM.

**Table 2.** Distribution of SNPs into genotype classes.

| Type | Parent 1 | Parent 2 | P1 dosage | P2 dosage | SNP initial number | Mapped SNP |
| --- | --- | --- | --- | --- | --- | --- |
| Null (N) | AAAA | BBBB | 4 | 0 | 8 | 0 |
| Simplex (S) | AAAA | AAAB | 4 | 3 | 1,332 | 591 |
| Simplex (S) | ABBB | BBBB | 1 | 0 |  |  |
| Triplex (T) | AAAA | ABBB | 4 | 1 | 126 | 0 |
| Triplex (T) | AAAB | BBBB | 3 | 0 |  |  |
| Duplex (D) | AAAA | AABB | 4 | 2 | 720 | 172 |
| Duplex (D) | AABB | BBBB | 2 | 0 |  |  |
| Double-simplex (SS) | AAAB | AAAB | 3 | 3 | 1,013 | 78 |
| Double-simplex (SS) | ABBB | ABBB | 1 | 1 |  |  |
| X-double-simplex (XSS) | AAAB | ABBB | 3 | 1 | 65 | 8 |
| Simplex-duplex (SD) | AAAB | AABB | 3 | 2 | 11 | 0 |
| Duplex-simplex (DS) | AABB | ABBB | 2 | 1 | 773 | 116 |
| Double-duplex (DD) | AABB | AABB | 2 | 2 | 192 | 35 |
| Total |  |  |  |  | 4,240 | 1,000 |

In Figure 3, the possible genotype configuration of each marker is indicated by the marker name prefix and the different colors. For example, markers with the prefix S and the color black represent the simplex configuration (Table 2). Four homologous groups showed almost all configurations (HG1, HG3, HG5 and HG9), and simplex markers (S) were the most frequent. The X-double-simplex (XSS) configuration was less frequent and appeared in only HG3 and HG5 (Table 2; Figure 3).

### Phenotypic Analysis and QTL Mapping

The selected VCOV matrix for ***G***_*a*_ based on AIC was CS-Het and/or US, in which both account for genetic correlation and heterogeneous variance across years. Considering the SIC, the selected VCOV matrix for ***G***_*a*_ was CS, in which the genetic correlation and homogeneous variance across years are considered, in addition to requiring estimation of a lower number of parameters than the previous VCOV matrices. As the difference in SIC values between CS and CS-Het/US (18880.37 – 18877.39 = 2.98) was greater than the respective AIC difference (18848.03 – 18845.13 = 2.9), the CS matrix was used for ***G***_*a*_ (Table 3).

**Table 3.** Values of AIC and SIC for ***G***_***a***_ matrix, considering different VCOV structures.

| $G_a$ matrix | Df | AIC | SIC | logREML | |
| --- | --- | --- | --- | --- | --- |
| ID | 4 | 18868.02 | 18891.51 | -9430.010 |  |
| DIAG | 5 | 18866.02 | 18895.41 | -9428.021 |  |
| CS | 5 | 18848.03 | **18877.39** | -9419.014 | Selected |
| CS-Het | 6 | **18845.13** | 18880.37 | -9416.565 |  |
| FA1 | 7 | 18847.13 | 18888.24 | -9416.565 |  |
| US | 6 | **18845.13** | 18880.37 | -9416.565 |  |

Predicted genotypic values for spittlebug (*Notozulia entreriana*) resistance in hybrids of the mapping population, genetic and residual variance components and trait heritability are shown in Table 4. The assumption of normality of the residuals can be visualized in Supplementary Figure 1.

**Table 4.**
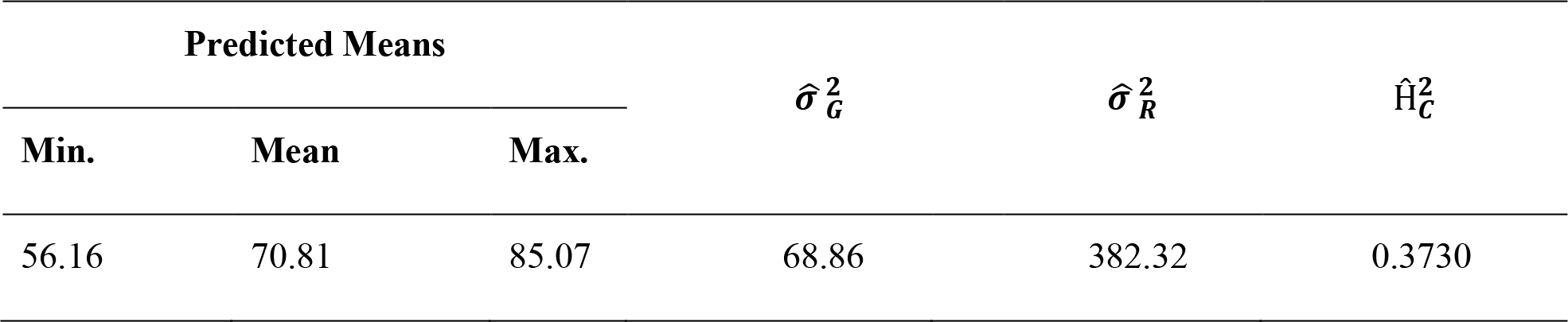
Summary of the predicted genotypic values, genetic 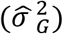 and residual 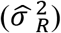 variance and heritability 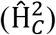 for spittlebug resistance.

Best linear unbiased prediction values were used to perform QTL mapping for spittlebug resistance by applying an IM model to the integrated genetic map. Using this method, three significant QTLs were identified for spittlebug resistance (Table 5) that were located in three homologous groups: HG1, HG2 and HG6. The obtained threshold values for LOD scores using 1,000 permutations ranged from 4.39 to 5.49 and were used to declare the presence of these QTLs. The percentages of phenotypic variation (*R*2) explained by the QTLs ranged from 4.66% to 6.24%. The male progenitor of the mapping population, *U. decumbens* D62 (cv. Basilisk), contributed with the resistance alleles for these three QTLs

**Table 5.**
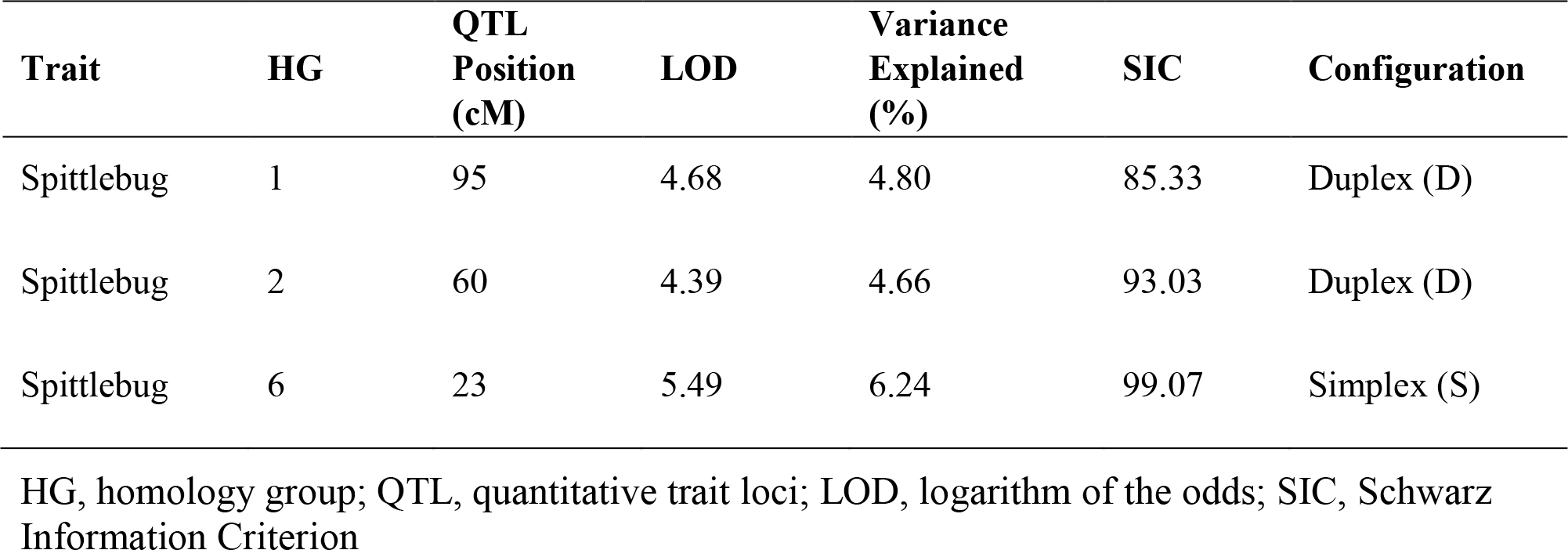
QTL information for the spittlebug resistance trait analyzed in the *U. decumbens* progeny.

| Trait | HG | QTL<br>Position<br>(cM) | LOD | Variance<br>Explained<br>(%) | SIC | Configuration |
| --- | --- | --- | --- | --- | --- | --- |
| Spittlebug | 1 | 95 | 4.68 | 4.80 | 85.33 | Duplex (D) |
| Spittlebug | 2 | 60 | 4.39 | 4.66 | 93.03 | Duplex (D) |
| Spittlebug | 6 | 23 | 5.49 | 6.24 | 99.07 | Simplex (S) |

HG, homology group; QTL, quantitative trait loci; LOD, logarithm of the odds; SIC, Schwarz Information Criterion

In homology group 1, the QTL had a LOD score of 4.68 and explained 4.80% of the phenotypic variation for spittlebug resistance. This LOD score was above the upper 95% LOD permutations threshold of 3.11. This first QTL was located at position 95 cM (Figure 3). Analyses of different simple genetic models were performed with TetraploidSNPMap to determine the best simple fitting model for the studied trait. For this QTL, the best model was a duplex genotype (AAAA × BBAA), with the allele B present on parent 2 associated with spittlebug resistance. This model had the lowest SIC (SIC = 85.34) compared with the full model (SIC = 102.99) (Table 5).

In homology group 2, the maximum LOD score was 4.39, and this QTL explained 4.66% of the phenotypic variation for spittlebug resistance. The LOD peak was located at position 60 cM (Figure 3), and its score was above the upper 95% LOD permutation threshold of 3.29 (Supplementary Figure 2). Again, the analysis with the simpler models estimated a duplex genotype (AAAA × BBAA) for this QTL, with the resistance allele present in parent 2. This model had the lowest SIC (SIC = 93.03) in comparison with the full model (SIC = 89.55) (Table 5).

The last detected QTL, in homology group 6, had the highest LOD score, 5.49, explained 6.24% of the phenotypic variation and was above the 95% LOD permutation upper threshold of 2.79 (Supplementary Figure 2). The QTL peak was located at 23 cM (Figure 3), and an analysis of different genetic models indicated that an allele explains the phenotypic variation. This QTL is linked to a simplex SNP (AAAA x ABAA), with the B allele present on parent 2 associated with spittlebug resistance. This model had the lowest SIC of 99.07 compared to the full model with SIC of 115.47 (Table 5).

### Search for Candidate Resistance Genes

Sequence similarity was detected in the regions of the markers adjacent to the mapped QTLs, with homologies for *Arabidopsis thaliana* and *Oryza sativa.* Functional annotation of these sequences showed possible candidate genes involved in the spittlebug resistance response (Supplementary Table 1) that are also present in the *U. decumbens* transcriptome (Salgado et al., 2017).

The first identified QTL, located in HG1, had two adjacent markers, D.1_40431816 and S.1_21570712, identified from the *S. viridis* genome reference. The region of this QTL showed similarity to pathogenesis-related gene 1 (*PR1*), leucine-rich repeat (LRR) family protein and the *WRKY11* gene, which are involved in pathogen defense (Supplementary Table 1).

In HG2, the region of the GBS-based adjacent markers (DS.2_39902899 and S.2_399022936) to QTL2 showed similarity to NBS-LRR and NB-ARC protein domains and with the pentatricopeptide repeat (PPR) superfamily proteins (Supplementary Table 1).

The region of the GBS-based markers (D.6_25761630 and S.6_35357677) adjacent to QTL3 in the HG6 showed similarity to a WRKY DNA-binding protein, a kinase superfamily protein, an F-box family protein, NB-ARC protein domains and kinase superfamily proteins. This QTL region also showed similarity to the thaumatin-like protein superfamily (TLP), which is classified within the pathogenesis-related (PR) protein family, to LRR protein domains and to F-box family proteins (Supplementary Table 1).

## 4 Discussion

The development of GBS technology enables the identification of thousands of SNP markers across the genome at relatively low cost. This advance represents an evolution in genetic studies of polyploids species, since a larger part of the genome can be represented, favoring approaches such as the construction of genetic maps. The map reported here was developed using an intraspecific progeny of *U. decumbens* of Embrapa Beef Cattle and includes GBS-based markers with autotetraploid allele dosages, an approach never previously employed in a forage grass species. Although not completely elucidated, some evidence suggests that *U. decumbens* is a segmental allopolyploid (Mendes-Bonato et al., 2001; Worthington et al., 2016), considered to have arisen from hybridization between very closely related species (Stebbins, 1947).

Our molecular markers fit a tetrasomic segregation model (Supplementary Figure 3), probably because autotetraploids as well as segmental allotetraploids, do not behave as diploids with respect to the mode of inheritance (Bourke et al., 2018). Another explanation is the fact that one of the parents of the mapping population was tetraploidized artificially and therefore this parent behaves as autotetraploid. We choose to use all possible SNP configurations to add even more genetic information to genetic map. Our map provides important genomic information, such as the detection of QTLs involved in spittlebug (*N. entreriana* Berg) resistance, a relevant trait for breeding programs in the *Urochloa* genus.

We used the genome of *S. viridis*, a member of the Poaceae, or grass, family, to align the tags and discover GBS-based markers. This alignment was sufficient to identify a relevant number (4,240) of high-quality SNPs, with allele dosage estimated. In previous analyses, we tested other reference genomes for tag alignment, but the *S. viridis* genome alignment produced the best results in terms of the number of markers identified (Supplementary Table 2). The sequences obtained herein by the GBS libraries are dynamic, allowing the raw data to be reanalyzed as bioinformatics techniques evolve (Poland and Rife, 2012), as well as when some tetraploid *Urochloa* spp. genome is sequenced. Therefore, our sequences may be reanalyzed in the future to obtain a higher alignment rate, as well as to identify a larger number of markers.

Worthington et al. (2016) detected 3,912 SNPs markers after filtering using the UNEAK (Universal Network-Enabled Analysis Kit) pipeline (Lu et al., 2013), which is similar to the number of markers that we detected using the *S. viridis* genome as reference. In a previous analysis, we tested the UNEAK pipeline for SNP identification in our intraspecific progeny, but this method detected fewer markers (1,210) compared with alignment to the genome reference (4,240). Most likely, the cross between two different species (*U. decumbens* × *U. ruziziensis*) favored the discovery of more SNPs through the UNEAK pipeline by Worthington et al. (2016), in addition to the different methodology used for GBS library construction.

Previous polyploid genetic studies have been developed with markers scored as presence/absence of alleles, but this method does not make use of all the information available in the SNP at multiple doses. The GBS data obtained here were sufficient in terms of read depth to call allele dosage in 9,345 markers from the *S. viridis* genome using SuperMASSA software (Serang et al., 2012; Pereira et al., submitted) (Supplementary Table 2). This methodology provided the distribution of alleles in the progeny and the relative intensities of each allele, increasing the amount of genetic information obtained (Serang et al., 2012; Garcia et al., 2013; Pereira et al., submitted).

After analysis using SuperMASSA software (Serang et al., 2012) and subsequent filtering steps, were obtained 4,240 SNPs that represented all possible configurations for an autotetraploid species.

Approximately 55% (2,345) of the SNPs markers identified from the *S. viridis* genome alignment followed the parental genotype configurations of simplex (AAAA x AAAB/ABBB x BBBB) and double-simplex (AAAB × AAAB/ABBB × ABBB). Slightly less than 1% of the identified markers followed the nulliplex (AAAA × BBBB) and simplex-duplex (AAAB × AABB) configurations (Table 2). These results were satisfactory and relevant for genetic studies with a tetraploid species, but a GBS library construction methodology with more than one restriction enzyme might provide a larger read depth and, accordingly, increase the accuracy of allele dosage estimation and the number of markers detected.

The 4,240 high-quality SNPs markers were inserted in TetraploidSNPMap software (Hackett et al., 2017), but only 1,515 showed significance in the χ^2^ goodness-of-fit statistic, and 1,000 were placed in the linkage map (Table 1). These 1,000 SNPs markers were distributed across nine homology groups, with a total length of 1,335.09 cM and an average map density of 1.33 cM (Table 1; Figure 3), reflecting the molecular source and chosen mapping methodology. Comparatively, Thaikua et al. (2016) and Vigna et al. (2016) presented the largest maps, with 1,702.82 cM and 1,423.2 cM of total size, respectively. Although small, this length difference may be due to the smaller number of non-NGS markers used (89 and 64, respectively) and the exclusive usage of SDMs for mapping. In addition to having a smaller length, our genetic map has a higher marker density than these others, reflecting the ability of the GBS methodology to allow the simultaneous detection of thousands of markers.

In the Worthington et al. (2016) study, genetic maps for an interspecific progeny resulting from a cross between *U. decumbens* × *U. ruziziensis* was built with GBS-based markers, and 1,916 SDA (single-dose allele) markers were distributed across the parental linkage maps. The different mapping method used in our study may have influenced the number of markers linked in the map. However, although our map has a lower marker density, it contain SNP markers with different allele dosage, adding more genetic information to the map and enabling more robust genetic analyses (Garcia et al., 2013).

In our study, the density of markers per HG ranged from 63 (HG6) to 187 (HG9) (Table 1). The greater density of markers for certain HGs may correspond to a greater recombination frequency. On the other hand, less saturated groups may have fewer SNP in these regions and/or correspond to highly homozygous regions that have lower recombination frequencies (Bai et al., 2016).

The largest distance between adjacent markers, 12.4 cM, was observed in homology group eight (Figure 3), one of the groups with the fewest mapped markers. Compared with other linkage map studies, our map has low average marker intervals in all HGs (Figures 2 and 3). Nevertheless, substantial intervals are common and even expected using the GBS technique, mainly due to centromeric regions, which are not reached with this methodology (Conson et al., 2018).

Additionally, it is possible that the number of hybrids in the mapping population was not sufficient to observe recombination in these intervals (Ma et al., 2014) or that these intervals represent regions that were not captured with the methodology used. One possible method to fill these distances is to use different combinations of enzymes to enlarge the sequencing pools and thereby enable the capture of markers in other genomic regions. Furthermore, we mapped only markers identified by aligning with the *S. viridis* genome. Therefore, these observed intervals may contain markers that are exclusive to the *U. decumbens* genome and thus cannot be represented with the methodology used.

Therefore, advances in the assembly of complex polyploid genomes are expected to enable the use of the full signalgrass genome as a reference for future studies (Margarido and Heckerman, 2015).

Of the 1000 mapped SNPs, 59% (591) followed the simplex parental genotype configuration (Table 2; Figure 3). This configuration was also the most common in other genetic mapping studies (Hackett et al., 2013; da Silva et al., 2017), probably because it is the most common in plant genomes and is easily scored. The least common configuration was X-double-simplex (XSS), followed by double-duplex (DD) (Table 2), which is the most complete configuration because it allows the representation of all types of segregation for autotetraploid species. Overall, the map displays most of the possible SNP configurations, and these data significantly increase the information about each locus and provide several advantages for genetic analysis (Garcia et al., 2013). When building genetic maps for polyploid species, the use of markers with allele dosage information allows the development of robust genetic maps that allow the detection of QTLs (Hackett et al., 2013; da Silva et al., 2017).

Until recently, no intraspecific progeny have been available for *U. decumbens*; therefore, no QTL mapping studies of this species have been performed to date. The identification of genes/loci underlying spittlebug (*N. entreriana*) resistance is critical for forage grass molecular breeding programs because these pests severely damage pastures (Valle et al., 2009). In our study, QTL mapping for spittlebug resistance was performed by applying the statistical model IM for the tetraploid progeny (Hackett et al., 2017). Although more accurate models are available, these models have not been extended to polyploidy and therefore do not use molecular markers with allele dosage information.

For the multi-allelic QTL analysis, we performed a permutation test to obtain the threshold for declaring significant QTLs. This analysis allowed the identification of three significant QTLs which explained between 4.66% and 6.24% of the phenotypic variation (Table 5). The identification of a small number of QTLs with small effects reflects the median heritability found for spittlebug resistance (*HC*^*2*^ = 0.37). Spittlebug resistance in forage grasses is probably not genetically complex but likely involves more than a single major resistance gene (Miles et al., 1995). Therefore, values of phenotypic variation and the number of QTLs detected in our analysis may be mainly due the low genetic variability for spittlebug resistance in the analyzed progeny, since the parents do not differ greatly for this trait. Moreover, the methodology used for QTL detection and/or map saturation may have influenced these results.

Although *U. decumbens* is considered a susceptible species to the pasture spittlebug, our results showed that the segregation of different alleles in each tetraploid parent contributed to the observed polymorphisms in the progeny. Therefore, our results can be useful for breeders to identify the alleles from the male parent that contribute to increased values of this phenotypic trait, consequently allowing genotypes that are more resistant to the development and survival of spittlebugs to be found more rapidly.

QTLs with greater effects on spittlebug resistance are necessary and more efficient for use in breeding programs, but our results represent a great gain as the only genetic architecture study using an *U. decumbens* intraspecific progeny. Therefore, the detection of these QTLs is the first step in the advancement of genomic studies involving spittlebug resistance. The annotation of sequences that originated markers with mapped QTLs is important to identify candidate genes involved in spittlebug resistance in *U. decumbens*. Thus, the regions that gave rise to these QTLs should be more thoroughly evaluated to identify more genetic information for future applications in the marker-assisted selection in signalgrass breeding programs.

*N. entreriana* spittlebug nymphs constantly suck the sap of roots, causing yellowing of the plant, and the saliva of the adults induces phytotoxemia that causes plant death (Valério and Nakano, 1992; Gusmão et al., 2016). Thus, to respond mechanical damage due to these insect attacks, plants possess multiple molecular defense mechanisms (War et al., 2012). Based on an investigative and comparative analysis, the QTL regions of this study showed similarity to genes containing protein domains reported to be involved in disease resistance and defense responses to pathogens that attack plants (Supplementary Table 1).

For spittlebug resistance trait, we can highlight the similarity of the region of the QTL1 with pathogenesis-related gene 1 (*PR1*). PR proteins are induced in response to attack by various types of pathogens, being that the *PR1* gene is involved in biological processes related to defense responses and systemic acquired resistance (Metzler et al., 1991; Uknes et al., 1992; Van Loon and Van Strien, 1999; Van Loon et al., 2006; Ali et al., 2018). This region also showed similarity to the *WRKY11* gene, which acts as a negative regulator of basal resistance to bacterial pathogens (Journot-Catalino et al., 2006) and plays a crucial role in cellular defense responses and defense-related gene transcriptional regulation (Jiang et al., 2016) (Supplementary Table 1).

For QTL3, we can highlight the similarity of the region to the adjacent markers with WRKY DNA binding domains. These transcription factors are involved in plant defense responses (Du and Chen, 2000). Moreover, F-box and WRKY DNA binding domains have orthologs in *A. thaliana*, with genes that develop a function in plant defense. This QTL region also showed similarity to TLPs, which play a variety of roles including plant protection against pathogen attacks (Liu et al., 2010; Mukherjee et al., 2010; Misra et al., 2016).

The regions of the markers adjacent to the mapped QTLs 2 and 3 showed similarity to NBS-LRR protein domains, which are widely distributed in plants and thought to respond to pathogen attacks, including viruses, bacteria, fungi, and even insects (Song et al., 2017). NBS-LRRs are the most prevalent class of resistance proteins (Gupta et al., 2012). F-box proteins form one of the largest multigene superfamilies and control many important biological functions such as pathogen resistance (Lechner et al., 2006), and the PPR superfamily proteins (Barkan and Small, 2014) have been implicated in plant defense mechanisms. Moreover, the NB-ARC protein domain is a functional ATPase domain containing a nucleotide-binding site that is proposed to regulate plant disease resistance (Van Ooijen et al., 2008).

Although we used the *S. viridis* genome as a reference to investigate the marker regions adjacent to the identified QTLs, we selected only genes found in the *U. decumbens* transcriptome (Salgado et al., 2017) to lend greater reliability to our results. Therefore, our results provide a starting point, indicating possible candidate genes involved in spittlebug resistance in signalgrass. However further studies should be conducted to validate these genomic regions as well as the role of the candidate genes in signalgrass and their effects on phenotypic expression.

We can conclude that this recent opportunity to analyze a *U. decumbens* full-sib progeny has opened new genetic molecular perspectives for this and related species. The genetic map presented herein includes important approaches such as the estimation of tetraploid dosage of each molecular SNP and greatly increases genetic information and represents an important evolution for polyploid studies (Bourke et al., 2016). In addition, the map allowed the identification of genomic regions related to spittlebug (*N. entreriana*) resistance, the main insect that attacks forage grasses, providing new insights about this trait. Furthermore, the molecular data developed here are dynamic and can be applied in future studies, such as genome assembly and other QTL analyses in *U. decumbens*.

Therefore, our results are the first step towards possible marker-assisted selection (MAS) and genomic selection (GS), a subject of great interest in genetic improvement programs for economically important plant species.

## Author Contributions

LC, SB, CV, JV, FZ and AS conceived and designed the experiments. RF, LC, SB, CV, JV and FZ conducted the experiments and collected data. RF, LL and AG analyzed the data. RF, LL and AS wrote the manuscript. All authors read and approved the manuscript.

## Acknowledgments

We would like to acknowledge the Brazilian Agricultural Research Corporation (Embrapa Beef Cattle) for providing the *Urochloa decumbens* progeny used in this study. We thank the Aline da Costa Lima Moraes for the assistance in the GBS library construction and performing DNA sequencing.

## Supplementary Figure and Tables Legends

**Supplementary Figure 1.**Residual plots.

**Supplementary Figure 2.**Interval mapping (IM) for spittlebug resistance from the *U. decumbens* mapping population in chromosomes 1 (A), 2 (B) and 6 (C). Dotted lines indicate the LOD thresholds of 90% and 95% obtained after the permutation tests.

**Supplementary Figure 3.**Theoretical distribution of genotypes in the mapping population and the distribution of individuals assigned to each genotype. The genotype distributions are nearly identical to the theoretical distribution in a F1 population.

**Supplementary Table 1.**Functional description of the sequences found in the regions of the markers adjacent to mapped *Urochloa decumbens* QTLs for spittlebug resistance based on similarity to the *Setaria viridis* genome.

**Supplementary Table 2.**Number of markers identified in the alignment with five reference genomes using Tassel-GBS pipeline after allele dosage estimation with SuperMASSA software and R data filtration analysis.

## References

Akaike, H. (1974). A new look at the statistical model identification. IEEE Trans. Automat. Contr. 19, 716–723.

Ali, S., Mir, Z.A., Bhat, J.A., Tyagi, A., Chandrashekar, N., Yadav, P., et al. (2018). Isolation and characterization of systemic acquired resistance marker gene PR1 and its promoter from Brassica juncea. 3 Biotech 8, 10. doi: 10.1007/s13205-017-1027-8

Almeida, M.C.D.C., Chiari, L., Jank, L., and Valle, C.B.D. (2011). Diversidade genética molecular entre cultivares e híbridos de Brachiaria spp. e Panicum maximum. Ciênc. Rural 41, 1998–2003.

Associação Brasileira das Indústrias Exportadoras de Carne (ABIEC) (2016). Perfil da Pecuária do Brasil: Relatório Anual. Available at: http://abiec.siteoficial.ws/images/upload/sumario-pt-010217.pdf

Bai, Z.-Y., Han, X.-K., Liu, X.-J., Li, Q.-Q., and Li, J.-L. (2016). Construction of a high-density genetic map and QTL mapping for pearl quality-related traits in Hyriopsis cumingii. Sci. Rep. 6, 32608. doi: 10.1038/srep32608

Barkan, A., and Small, I. (2014). Pentatricopeptide repeat proteins in plants. Annu. Rev. Plant Biol. 65, 415–442. doi: 10.1146/annurev-arplant-050213-040159

Bourke, P.M., Voorrips, R.E., Kranenburg, T., Jansen, J., Visser, R.G.F., and Maliepaard, C. (2016). Integrating haplotype-specific linkage maps in tetraploid species using SNP markers. Theor. Appl. Genet. 129, 2211–2226. doi: 10.1007/s00122-016-2768-1

Bourke, P.M., Voorrips, R.E., Visser, R.G.F., and Maliepaard, C. (2018). Tools for genetic studies in experimental populations of polyploids. Front. Plant Sci. 9, 513. doi:10.3389/fpls.2018.00513

Brien, C. (2016). ASRemlPlus: augments the use of ‘ASReml’ in fitting mixed models. R package version 2. 0–9.

Butler, D.G., Cullis, B.R., Gilmour, A.R., and Gogel, B.J. (2009). ASReml-R reference manual, release 3. Technical report. Brisbane, QLD: Queensland Department of Primary Industries.

Conson, A.R.O., Taniguti, C.H., Amadeu, R.R., Andreotti, I.A.A., de Souza, L.M., Dos Santos, L.H.B., et al. (2018). High-resolution genetic map and QTL analysis of growth-related traits of Hevea brasiliensis cultivated under suboptimal temperature and humidity conditions. Front. Plant Sci. 9, 1255. doi: 10.3389/fpls.2018.00513

Cullis, B.R., Smith, A.B., and Coombes, N.E. (2006). On the design of early generation variety trials with correlated data. J. Agric. Biol. Environ. Stat. 11, 381–393. doi: 10.1198/108571106X154443

Da Silva, W.L., Ingram, J., Hackett, C.A., Coombs, J.J., Douches, D., Bryan, G.J., et al. (2017). Mapping loci that control tuber and foliar symptoms caused by PVY in Autotetraploid potato (Solanum tuberosum L.). G3 7, 3587–3595. doi: 10.1534/g3.117.300264

Du, L., and Chen, Z. (2000). Identification of genes encoding receptor-like protein kinases as possible targets of pathogen- and salicylic acid-induced WRKY DNA-binding proteins in Arabidopsis. Plant J. 24, 837–847. doi: 10.1046/j.1365-313x.2000.00923.x

Ebina, M., Nakagawa, H., Yamamoto, T., Araya, H., Tsuruta, S.-i., Takahara, M., et al. (2005). Co-segregation of AFLP and RAPD markers to apospory in Guineagrass (Panicum maximum Jacq.). Grassl. Sci. 51, 71–78. doi: 10.1111/j.1744-697X.2005.00011.x

Elshire, R.J., Glaubitz, J.C., Sun, Q., Poland, J.A., Kawamoto, K., Buckler, E.S., et al. (2011). A robust, simple genotyping-by-sequencing (GBS) approach for high diversity species. PLoS One 6, e19379. doi: 10.1371/journal.pone.0019379

Ferreira, R.C.U., Cançado, L.J., Do Valle, C.B., Chiari, L., and De Souza, A.P. (2016). Microsatellite loci for Urochloa decumbens (Stapf) R.D. Webster and cross-amplification in other Urochloa species. BMC Res. Notes 9, 152. doi: 10.1186/s13104-016-1967-9

Garcia, A.A.F., Mollinari, M., Marconi, T.G., Serang, O.R., Silva, R.R., Vieira, M.L.C., et al. (2013). SNP genotyping allows an in-depth characterisation of the genome of sugarcane and other complex autopolyploids. Sci. Rep. 3, 3399. doi: 10.1038/srep03399

Gilmour, A.R., Gogel, B.J., Cullis, B.R., and Thompson, R. (2009). ASReml User Guide Release 3.0. Hempstead, NY: VSN Int. Ltd.

Glaubitz, J.C., Casstevens, T.M., Lu, F., Harriman, J., Elshire, R.J., Sun, Q., et al. (2014). TASSEL-GBS: a high capacity genotyping by sequencing analysis pipeline. PLoS One 9, e90346. doi: 10.1371/journal.pone.0090346

Goodstein, D.M., Shu, S., Howson, R., Neupane, R., Hayes, R.D., Fazo, J., et al. (2012). Phytozome: a comparative platform for green plant genomics. Nucl. Acids Res. 40, D1178–D1186. doi: 10.1093/nar/gkr944

Gupta, S.K., Rai, A.K., Kanwar, S.S., and Sharma, T.R. (2012). Comparative analysis of zinc finger proteins involved in plant disease resistance. PLoS One 7, e42578. doi: 10.1371/journal.pone.0042578

Gusmão, M.R., Valério, J.R., Matta, F.P., Souza, F.H.D., Vigna, B.B.Z., Fávero, A.P., et al. (2016). Warm-season (C4) turfgrass genotypes resistant to spittlebugs (Hemiptera: Cercopidae). J. Econ. Entomol. 109, 1914–1921. doi: 10.1093/jee/tow135

Hackett, C.A., Boskamp, B., Vogogias, A., Preedy, K.F., and Milne, I. (2017). TetraploidSNPMap: software for linkage analysis and QTL mapping in autotetraploid populations using SNP dosage data. J. Hered. 108, 438–442. doi: 10.1093/jhered/esx022

Hackett, C.A., Bradshaw, J.E., and Bryan, G.J. (2014). QTL mapping in autotetraploids using SNP dosage information. Theor. Appl. Genet. 127, 1885–1904. doi: 10.1007/s00122-014-2347-2

Hackett, C.A., McLean, K., and Bryan, G.J. (2013). Linkage analysis and QTL mapping using SNP dosage data in a tetraploid potato mapping population. PLoS One 8, e63939. doi: 10.1371/journal.pone.0063939

Heffelfinger, C., Fragoso, C.A., Moreno, M.A., Overton, J.D., Mottinger, J.P., Zhao, H., et al. (2014). Flexible and scalable genotyping-by-sequencing strategies for population studies. BMC Genom. 15, 979. doi: 10.1186/1471-2164-15-979

Jank, L., Valle, C.B., and Resende, R.M.S. (2011). Breeding tropical forages. Crop Breed. Appl. Biotechnol. 11, 27–34. doi: 10.1590/S1984-70332011000500005

Jiang, C.-H., Huang, Z.-Y., Xie, P., Gu, C., Li, K., Wang, D.-C., et al. (2016). Transcription factors WRKY70 and WRKY11 served as regulators in rhizobacterium Bacillus cereus AR156- induced systemic resistance to Pseudomonas syringae pv.tomato DC3000 in Arabidopsis. J. Exp. Bot. 67, 157–174. doi: 10.1093/jxb/erv445

Journot-Catalino, N., Somssich, I.E., Roby, D., and Kroj, T. (2006). The transcription factors WRKY11 and WRKY17 act as negative regulators of basal resistance in Arabidopsis thaliana. Plant Cell Online 18, 3289–3302. doi: 10.1105/tpc.106.044149

Langmead, B., and Salzberg, S.L. (2012). Fast gapped-read alignment with Bowtie 2. Nat. Methods 9, 357–359. doi: 10.1038/nmeth.1923

Lapointe, S.L., Serrano, M.S., Arango, G.L., Sotelo, G., and Cordoba, F. (1992). Antibiosis to spittlebugs (Homoptera: Cercopidae) in accessions of Brachiaria spp. J. Econ. Entomol. 85, 1485–1490. doi: 10.1093/jee/85.4.1485

Lechner, E., Achard, P., Vansiri, A., Potuschak, T., and Genschik, P. (2006). F-box proteins everywhere. Curr. Opin. Plant Biol. 9, 631–638. doi: 10.1016/j.pbi.2006.09.003

Liu, J.-J., Sturrock, R., and Ekramoddoullah, A.K.M. (2010). The superfamily of thaumatin-like proteins: its origin, evolution, and expression towards biological function. Plant Cell Rep. 29, 419–436. doi: 10.1007/s00299-010-0826-8

Lu, F., Lipka, A.E., Glaubitz, J., Elshire, R., Cherney, J.H., Casler, M.D., et al. (2013). Switchgrass genomic diversity, ploidy, and evolution: novel insights from a network-based SNP discovery protocol. PLoS Genet. 9, e1003215. doi: 10.1371/journal.pgen.1003215

Ma, J.-Q., Yao, M.-Z., Ma, C.-L., Wang, X.-C., Jin, J.-Q., Wang, X.-M., et al. (2014). Construction of a SSR-based genetic map and identification of QTLs for catechins content in tea plant (Camellia sinensis). PLoS One 9, e93131. doi: 10.1371/journal.pone.0093131

Margarido, G.R.A., and Heckerman, D. (2015). ConPADE: genome assembly ploidy estimation from next-generation sequencing data. PLoS Comput. Biol. 11, e1004229. doi: 10.1371/journal.pcbi.1004229

Mendes-Bonato, A.B., Pagliarini, M.S., Da Silva, N., and Do Valle, C.B. (2001). Meiotic instability in invader plants of signal grass Brachiaria decumbens Stapf (Gramineae). Acta Sci. 23, 619–625.

Metzler, M.C., Cutt, J.R., and Klessig, D.F. (1991). Isolation and characterization of a gene encoding a PR-1-like protein from Arabidopsis thaliana. Plant Physiol. 96, 346–348.

Miles, J.W., Lapointe, S.L., Escandén, M.L., and Sotelo, G. (1995). Inheritance of resistance to spittle bug (Homoptera: Cercopidae) in inter specific Brachiaria spp. hybrids. J. Econ. Entomol. 88, 1477–1481. doi: 10.1093/jee/88.5.1477

Misra, R.C., Sandeep , Kamthan, M., Kumar, S., and Ghosh, S. (2016). A thaumatin-like protein of Ocimum basilicum confers tolerance to fungal pathogen and abiotic stress in transgenic Arabidopsis. Sci. Rep. 6, 25340. doi: 10.1038/srep25340

Mollinari, M., and Serang, O. (2015). Quantitative SNP genotyping of polyploids with MassARRAY and other platforms. Methods Mol. Biol. 1245, 215–241. doi: 10.1007/978-1-4939-1966-6_17

Mukherjee, A.K., Carp, M.-J., Zuchman, R., Ziv, T., Horwitz, B.A., and Gepstein, S. (2010). Proteomics of the response of Arabidopsis thaliana to infection with Alternaria brassicicola. J. Proteom. 73, 709–720. doi: 10.1016/j.jprot.2009.10.005

Naumova, T.N., Hayward, M.D., and Wagenvoort, M. (1999). Apomixis and sexuality in diploid and tetraploid accessions of Brachiaria decumbens. Sex Plant Reprod. 12, 43–52.

Okada, M., Lanzatella, C., Saha, M.C., Bouton, J., Wu, R., and Tobias, C.M. (2010). Complete switchgrass genetic maps reveal subgenome collinearity, preferential pairing and multilocus interactions. Genetics 185, 745–760. 10.1534/genetics.110.113910

Patel, R.K., and Jain, M. (2012). NGS QC toolkit: a toolkit for quality control of next generation sequencing data. PLoS One 7, e30619. doi: 10.1371/journal.pone.0030619

Pereira, G.S., Garcia, A.A.F., and Margarido, G.R.A. A fully automated pipeline for quantitative genotype calling from next generation sequencing data in polyploids. Submitted.

Pessoa-Filho, M., Martins, A.M., and Ferreira, M.E. (2017). Molecular dating of phylogenetic divergence between Urochloa species based on complete chloroplast genomes. BMC Genom. 18, 516. doi: 10.1186/s12864-017-3904-2

Piepho, H. P., and Möhring, J. (2007). Computing heritability and selection response from unbalanced plant breeding trials. Genetics 177, 1881–1888. doi: 10.1534/genetics.107.074229

Poland, J.A., Brown, P.J., Sorrells, M.E., and Jannink, J.-L. (2012). Development of high-density genetic maps for barley and wheat using a novel two-enzyme genotyping-by-sequencing approach. PLoS One 7, e32253. doi: 10.1371/journal.pone.0032253

Poland, J.A., and Rife, T.W. (2012). Genotyping-by-sequencing for plant breeding and genetics. Plant Genome J. 5, 92–102. doi:10.3835/plantgenome2012.05.0005

Rajput, S.G., Santra, D.K., and Schnable, J. (2016). Mapping QTLs for morpho-agronomic traits in proso millet (Panicum miliaceum L.). Mol. Breed. 36, 37. doi: 10.1007/s11032-016-0460-4

Renvoize, S.A., Clayton, W.D., and Kabuye, C.H.S. (1996). “Morphology, taxonomy, and natural distribution of Brachiaria (Trin.). Griseb,” in Brachiaria: Biology, Agronomy, and Improvement, eds. J.W. Miles, B.L. Maass and C.B. Valle. (Cali, Colombia: Centro Internacional de Agricultura Tropical (CIAT), Empresa Brasileira de Pesquisa Agropecuaria (EMBRAPA)), 1–15.

Ripol, M.I., Churchill, G.A., da Silva, J.A.G., and Sorrells, M. (1999). Statistical aspects of genetic mapping in autopolyploids. Gene 235, 31–41.

Salgado, L.R., Lima, R., Santos, B.F.d., Shirakawa, K.T., Vilela, M.D.A., Almeida, N.F., et al. (2017). De novo RNA sequencing and analysis of the transcriptome of signalgrass (Urochloa decumbens) roots exposed to aluminum. Plant Growth Regul. 83, 157–170. doi: 10.1007/s10725-017-0291-2

Schwarz, G. (1978). Estimating the dimension of a model. Ann. Stat. 6, 461–464.

Serang, O., Mollinari, M., and Garcia, A.A.F. (2012). Efficient exact maximum a posteriori computation for Bayesian SNP genotyping in polyploids. PLoS One 7, e30906. doi: 10.1371/journal.pone.0030906

Shirasawa, K., Tanaka, M., Takahata, Y., Ma, D., Cao, Q., Liu, Q., et al. (2017). A high-density SNP genetic map consisting of a complete set of homologous groups in autohexaploid sweetpotato (Ipomoea batatas). Sci. Rep. 7, 44207. doi: 10.1038/srep44207

Simioni, C., and Valle, C.B. (2009). Chromosome duplication in Brachiaria (A. Rich.) stapf allows intraspecific crosses. Crop Breed. Appl. Biotechnol. 9, 328–334. doi: 10.12702/1984- 7033.v09n04a07

Song, H., Wang, P., Li, C., Han, S., Zhao, C., Xia, H., et al. (2017). Comparative analysis of NBS-LRR genes and their response to Aspergillus flavus in Arachis. PLoS One 12, e0171181. doi: 10.1371/journal.pone.0171181

Souza, J.S., Chiari, L., Simeão, R.M., de Mendonça Vilela, M., and Salgado, L.R. (2018). Development, validation and characterization of genic microsatellite markers in Urochloa species. Am. J. Plant Sci. 09, 281–295. doi: 10.4236/ajps.2018.92023

Stebbins, G. L. (1947). Types of polyploids: their classification and significance. Adv. Genet. 1, 403–429.

Stein, J., Pessino, S.C., Martínez, E.J., Rodriguez, M.P., Siena, L.A., Quarin, C.L., et al. (2007). A genetic map of tetraploid Paspalum notatum Flügge (bahiagrass) based on single-dose molecular markers. Mol. Breed. 20, 153–166. doi: 10.1007/s11032-007-9083-0

Thaikua, S., Ebina, M., Yamanaka, N., Shimoda, K., Suenaga, K., and Kawamoto, Y. (2016). Tightly clustered markers linked to an apospory-related gene region and quantitative trait loci mapping for agronomic traits in Brachiaria hybrids. Grassl. Sci. 62, 69–80. doi: 10.1111/grs.12115

Triviño, N.J., Perez, J.G., Recio, M.E., Ebina, M., Yamanaka, N., Tsuruta, S.-i., et al. (2017). Genetic diversity and population structure of species and breeding populations. Crop Sci. 57, 1–12. doi: 10.2135/cropsci2017.01.0045

Uknes, S., Mauch-Mani, B., Moyer, M., Potter, S., Williams, S., Dincher, S., et al. (1992). Acquired resistance in Arabidopsis. Plant Cell 4, 645–656. doi: 10.1105/tpc.4.6.645

Valério, J., and Nakano, O. (1988). Influência do adulto de Zulia entreriana (Berg, 1879) (Homoptera: Cercopidae) na produção e qualidade de Brachiaria decumbens. Pesqui. Agropecu. Bras. 23, 447–453.

Valério, J., and Nakano, O. (1992). Sintomatologia dos danos causados pelo adulto da cigarrinha Zulia entreriana (Berg, 1879) (Homoptera: Cercopidae) em Brachiaria decumbens Stapf. Ann. Soc. Entomol. Bras 21, 95–100.

Valério, J.R., Jeller, H., and Peixer, J. (1997). Seleçães do introduções do gênero Brachiaria (Griseb) resistentes à cigarrinha Zulia entreriana (Berg) (Homoptera: Cercopidae). An. Soc. Entomol. Bras. 26, 383–387.

Valle, C.B., Jank, L., and Resende, R.M.S. (2009). O melhoramento de forrageiras tropicais no Brasil. Rev. Ceres 56, 460–472.

Valle, C.B., Macedo, M.C.M., Euclides, V.P.B., Jank, L., and Resende, R. (2010). “Gênero Brachiaria,” in Plantas Forrageiras, eds. D.M.D.F. Martuscello and J. Azevedo. (Viçosa, Brasil: Editora UFV), 30–77.

Valle, C.B., Simioni, C., Resende, R.M.S., and Jank, L. (2008). “Melhoramento genético de Brachiaria,” in Melhoramento de Forrageiras Tropicais, eds. R.M.S. Resende, C.B. Valle and L. Jank. (Campo Grande, MS: Embrapa Gado de Corte), 13–54.

Van Loon, L.C., Rep, M., and Pieterse, C.M.J. (2006). Significance of inducible defense-related proteins in infected plants. Annu. Rev. Phytopathol. 44, 135–162. doi: 10.1146/annurev.phyto.44.070505.143425

Van Loon, L.C., and Van Strien, E.A. (1999). The families of pathogenesis-related proteins, their activities, and comparative analysis of PR-1 type proteins. Physiol. Mol. Plant Pathol. 55, 85–97.

Van Ooijen, G., Mayr, G., Kasiem, M.M.A., Albrecht, M., Cornelissen, B.J.C., and Takken, F.L.W. (2008). Structure–function analysis of the NB-ARC domain of plant disease resistance proteins. J. Exp. Bot. 59, 1383–1397. doi: 10.1093/jxb/ern045

Vigna, B.B.Z., Santos, J.C.S., Jungmann, L., do Valle, C.B., Mollinari, M., Pastina, M.M., et al. (2016). Evidence of allopolyploidy in Urochloa humidicola based on cytological analysis and genetic linkage mapping. PLoS One 11, e0153764. doi: 10.1371/journal.pone.0153764

Voorrips, R.E. (2002). MapChart: software for the graphical presentation of linkage maps and QTLs. J. Hered. 93, 77–78. doi: 10.1093/jhered/93.1.77

Voorrips, R.E., Gort, G., and Vosman, B. (2011). Genotype calling in tetraploid species from bi-allelic marker data using mixture models. BMC Bioinform. 12, 172. doi: 10.1186/1471-2105-12-172

War, A.R., Paulraj, M.G., Ahmad, T., Buhroo, A.A., Hussain, B., Ignacimuthu, S., et al. (2012). Mechanisms of plant defense against insect herbivores. Plant Signal Behav. 7, 1306–1320. doi: 10.4161/psb.21663

Worthington, M., Heffelfinger, C., Bernal, D., Quintero, C., Zapata, Y.P., Perez, J.G., et al. (2016). A parthenogenesis gene candidate and evidence for segmental allopolyploidy in apomictic Brachiaria decumbens. Genetics 203, 1117–1132. doi: 10.1534/genetics.116.190314

Wu, K.K., Burnquist, W., Sorrells, M.E., Tew, T.L., Moore, P.H., and Tanksley, S.D. (1992). The detection and estimation of linkage in polyploids using single-dose restriction fragments. Theor. Appl. Genet. 83, 294–300. doi: 10.1007/BF00224274

